# A biomimetic liver model recapitulating bio-physical properties and tumour stroma interactions in hepatocellular carcinoma

**DOI:** 10.1101/2020.04.30.069823

**Authors:** C. Calitz, N. Pavlovic, J. Rosenquist, C. Zagami, A. Samanta, F. Heindryckx

**Author notes:** Corresponding author: Femke Heindryckx. Email addresses of co-authors: Carlemi Calitz, Natasa Pavlovic, Jenny Rosenquist, Claudia Zagami, Ayan Samanta, Femke Heindryckx.

## Abstract

Hepatocellular carcinoma (HCC) is a primary liver tumor developing in the wake of chronic liver disease. Chronic liver disease and inflammation leads to a fibrotic environment actively supporting and driving hepatocarcinogenesis. Insight into hepatocarcinogenesis in terms of the interplay between the tumor stroma micro-environment and tumor cells is thus of considerable importance. Three-dimensional (3D) cell culture models are proposed as the missing link between current *in vitro* 2D cell culture models and *in vivo* animal models. Our aim was to design a novel 3D biomimetic HCC model with accompanying fibrotic stromal compartment and vasculature. Physiologically relevant hydrogels such as collagen and fibrinogen were incorporated to mimicking the bio-physical properties of the tumor ECM. In our model LX2 and HepG2 cells embedded in a hydrogel matrix were seeded onto the inverted insert membrane of a Transwell™ system. HUVEC cells were then seeded onto the opposite side of the membrane. Three formulations consisting of ECM-hydrogels embedded with cells were prepared and the bio-physical properties determined by rheology. Cell viability was determined by the AlamarBlue^®^ assay over 21-days. The effect of the chemotherapeutic drug doxorubicin was evaluated in both a 2D co-culture and our 3D model for a period of 72h. We show that this model is viable for 25-days and gives rise to metastatic tumor nodules after 17 days in culture. Rheology results show that bio-physical properties of a fibrotic, cirrhotic and HCC liver can be successfully mimicked. Overall, results indicate that this 3D model is more representative of the *in vivo* situation compared to traditional 2D cultures. Our 3D tumor model showed a decreased response to chemotherapeutics, mimicking drug resistance typically seen in HCC patients. This model could in future provide a valuable new platform to study multifocal HCC or to identify mechanisms that contribute to early stages of metastasis.

**SUMMARY:** A protocol for a novel 3D biomimetic HCC model with accompanying fibrotic stromal compartment and vasculature, to study endocrine and paracrine signaling in liver cancer. The model uses physiological relevant hydrogels in ratios mimicking the bio-physical properties of the stromal extracellular matrix, which is an active mediator of cellular interactions, tumor growth and metastasis.

## INTRODUCTION

Hepatocelluar carcinoma (HCC) comprises 90% of all primary liver cancers (Easl guidelines, 2018; Marquardt *et al*., 2015). With 810 000 deaths and 854 000 new cases reported annually it is currently ranked as the fifth most common cancer worldwide with one of the highest incidence of mortality (Easl guidelines, 2018). The development of HCC is predominantly attributed to inflammation associated with chronic liver diseases namely, viral hepatitis, chronic excessive alcohol intake, metabolic syndrome, obesity and diabetes. (Easl guidelines, 2018; Balogh *et al*., 2016; Perumpail *et al*., 2015). The inflammation associated with these pathological conditions results in hepatocyte injury and secretion of various cytokines that activate and recruit hepatic stellate cells and inflammatory cells to initiate fibrosis (Baglieri *et al*., 2019). Hepatic stellate cells are known for their key role in the initiation, progression and regression of hepatic fibrosis. Upon activation they differentiate into myofibroblast like cells with contractile, pro-inflammatory and pro-fibrinogenic properties (Arriazu *et al*., 2014; Malarkey *et al*.,2005; Moreira, 2007). The resulting fibrosis in turn causes the dysregulation of extracellular matrix remodeling enzyme activity, creating an environment characterized by an overall increased stiffness accompanied by the secretion of growth factors, which further contributes to HCC pathogenesis (Hernandes-Gea *et al*., 2013; Amicone & Marchetti, 2018). It is this continuous pathogenic feedback loop between hepatocytes and the stromal environment, which fuels cancer initiation, epithelial to mesenchymal transitions (EMT), metastatic potential, and altered drug response (Rawal *et al*., 2019; Yoo *et al*., 2017; Landry *et al*., 2018). Insight into hepatocarcinogenesis in terms of the interplay between the tumor and the tumor micro-environment is thus of considerable importance not only from a mechanistic but also from a treatment perspective.

Two-dimensional (2D) *in vitro* cell culture models are predominantly used by 80% of cancer cell biologists (Le *et al*., 2018). However, these models are not representative of the true tumor micro-environment, which affects chemotherapeutic responses (Lv *et al*., 2017; Hoarau-Véchot *et al*., 2018; Le *et al*., 2018). Currently 96% of chemotherapeutic drugs fail during clinical trials (Le *et al*., 2018). This high incidence in drug attrition rates can be attributed to the fact that available *in vitro* pre-screening models do not fully represent our current insight and understanding of HCC complexity and the microenvironment (Hoarau-Véchot *et al*., 2018). Conversely *in vivo* animal models present with compromised immune systems and discrepancies in interactions between the tumor and the microenvironment when compared to humans (Hoarau-Véchot *et al*., 2018; Khawar *et al*., 2018). On average only 8% of results obtained from animal studies can be reliably translated from the pre-clinical to the clinical setting (Hoarau-Véchot *et al*., 2018; Khawar *et al*., 2018). Thus, it is clear that the evaluation of HCC requires the development of an *in vitro* platform that effectively recapitulates the complexity of not only the tumor but also the microenvironment. A platform that can supplement current *in vitro* pre-clinical screening models and reduce the amount of *in vivo* animal studies (Malarkey *et al*., 2005; Le *et al*., 2018).

One such platform is advanced three-dimensional (3D) cell culture models. A multitude of these advanced 3D models to study HCC have emerged over the last decade and various reviews have been published. Available 3D models to study HCC include, multicellular spheroids, organoids, scaffold based models, hydrogels, microfluidics and bio-printing. Of these, multicellular spheroids are one of the best known models used in the study of tumor development. Spheroids are an inexpensive model with low technical difficulty while at the same time effectively mimicking *in vivo* tumor architecture (Elliott and Yuan, 2010; Nath and Devi, 2016; Zanoni *et al*., 2016). That multicellular spheroids have contributed to a wealth of information on HCC is undeniable (Bell *et al*., 2016; Jung *et al*., 2017; Khawar *et al*. 2018). However, standardized culture time is lacking as multicellular spheroids are kept in culture from between 7 to 48 days. Increased culture time is of considerable importance as, Eilenberger found that the differences in spheroid age profoundly influences Sorafinib diffusivity and toxicity (Eilenberger *et al*., 2019). While Wrzesinski and Fey found that 3D hepatocyte spheroids require 18 days to re-establish key physiological liver functions after trypsinization, and continues to exhibit stable functionality for up to 24 days after this recovery (Wrzesinski and Fey, 2013; Wrzesinski *et al*., 2013).

Some of the more advanced 3D HCC models includes the use of human decellularized liver scaffolds and bio-printed scaffolds. Mazza and colleagues created a natural 3D scaffold for HCC modelling using decellularized human livers not suitable for transplantation. These natural scaffolds could successfully be repopulated for 21 days with a co-culture of hepatic stellate and hepatoblastoma cells, while maintaining the expression of key extracellular matrix components such as collagen type I, III, IV and fibronectin. Other than disease modelling this model also offers the advantage of functional organ transplantation and pre-clinical drug and toxicity screening (Mazza *et al*., 2015). With the advances in 3D bio-printing, 3D extracellular matrix scaffolds can now also be bio-printed. Ma and colleagues bio-printed extracellular matrix scaffolds with variable mechanical properties and biomimetic microarchitecture using hydrogels engineer from decellularized extracellular matrix (Ma *et al*., 2018). Undoubtedly these are all excellent 3D HCC models. However, unavailability of human livers and the cost involved in acquiring the necessary equipment and materials places these models at a disadvantage. Additionally these methods are all technically advanced requiring extensive training that may not be readily available to all researchers.

Based on the complexity of HCC and currently available 3D models we endeavored to develop an all-encompassing 3D HCC model. We aimed for a model capable of recapitulating both the premalignant and tumor microenvironment by incorporating some of the key players in the development of HCC. These include endothelial cells, hepatic stellate cells and malignant hepatocytes, grown in a microenvironment composed of physiologically relevant hydrogels. With the chosen hydrogels, collagen type I and fibrinogen, incorporated in ratios comparable to bio-physical changes seen in liver stiffness during the initiation and progression of HCC. Additionally we aimed for a model that could be kept in culture for a prolonged time period. We envisioned a modular, cost effective model that can be setup with basic equipment, minimal training and experience, and readily available materials.

## PROTOCOL

### 1. Preparation of Fibrinogen stock solution

1.1 Prepare a 1 M calcium chloride (CaCl_2_) stock solution, by weighing 2.21 g CaCl_2_ and adding it to 20 mL distilled water (dH_2_O). Stock solution can be stored at room temperature.
1.2 Prepare a 20 mL aprotinin stock solution (1218.75 KIU/mL) by weighing 5 mg of aprotinin and adding it to 20 mL dH_2_O. Aliquot stock solution into 1 mL aliquots and store at -20 °C.
1.3 Prepare a 10 mL fibrinogen stock solution, add 7.849 mL phosphate buffered saline (PBS), 2.051 mL aprotinin stock (1218.75 KIU/mL) for a final concentration of 250 KIU/mL and 100 µL CaCl_2_ (1M) for a final concentration of 10 mM, into a 50 mL tube.
1.4 For a 10 ml, 80 mg/ml fibrinogen stock solution, weigh 800 mg fibrinogen.
1.5 For the stock solution to contain 2% w/v sodium chloride (NaCl) weigh 200 mg NaCl.
1.6 Gently add the fibrinogen and NaCl to the 50 mL tube containing the PBS, aprotinin and CaCl_2_. During this step it is very important that the solution is not stirred or shaken vigorously as this will result in fibrinogen gelling and lumps forming in the solution. Instead it is recommended to add fibrinogen and NaCl in increments when producing larger volumes of the stock solution and to place it on a shaker at a low setting of 300 r.p.m.
1.7 Once the solution has dissolved it can be filtered using a .22 micron syringe filter. Important, do not autoclave the fibrinogen solution as this will destroy the fibrinogen.

NOTE: This part of the protocol can take between 2 to 5 hours depending on the amount of stock solution, this time should be taken into consideration during the experimental setup.

### 2. Coating inserts with collagen

2.1 In a laminar flow hood, using sterilized tweezers, remove inserts from the plate and place inverted onto the lid of the plate.
2.2 Prepare 100 mL of a 20 mM acetic acid solution by adding 115 µL acetic acid to 25 mL dH_2_O and adjusting to a final volume of 100 mL with dH_2_O. The solution can then be filtered using a .22 micron syringe filter. Stock solution can be stored at room temperature.
2.3 Prepare 2 mL of a 100 µg/mL collagen solution from a 5 mg/mL collagen stock solution by adding 40 µL collagen to 1.060 mL acetic acid (20 mM).
2.4 Coat inserts with the 100 µg/mL collagen solution by pipetting 100 µL of the solution onto each insert. Let inserts air dry within the laminar flow hood.
2.5 Once inserts have dried wash each insert three times with 200 µL PBS. Let inserts air dry within the laminar flow hood.
2.6 Add a custom 3D printed spacer over the insert, this will be necessary once the gels are hanging from the insert to prevent them from touching the bottom well of the plate.
2.7 Cover the inverted inserts with the bottom part of the plate and place into the incubator until cells are embedded in the hydrogels and ready to be seeded

CAUTION: Acetic acid is toxic to cells and inserts should be washed thoroughly with PBS.

### 3. Seeding cells embedded in hydrogels onto inserts

The following table provides a description of the three formulations with varying concentrations fibrinogen that will be prepared. Formulation one corresponds to the liver during HCC, two during cirrhosis and three during the onset of fibrosis.

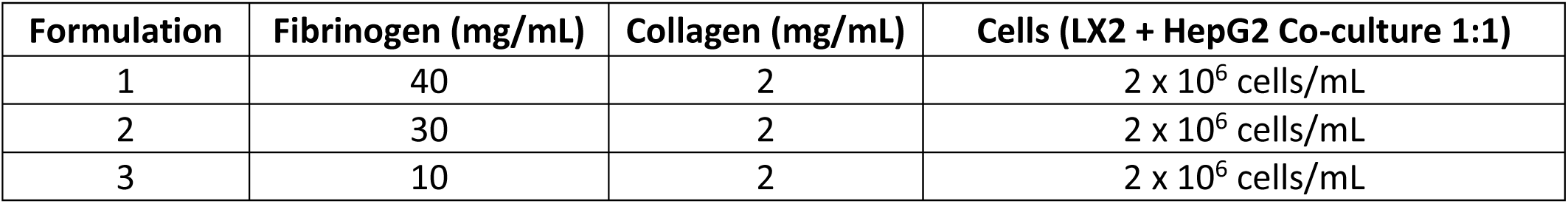

3.1 Prepare 1 M sodium hydroxide stock solution by adding 3.99 g NaOH to 100 mL dH_2_O. The solution can then be filtered using a .22 micron syringe filter. Stock solution can be kept at room temperature.
3.2 Collect ice and place the collagen stock 5 mg/mL and 10 mL 1M NaOH on ice.
3.3 In a water bath, preheat 50 mL PBS, 15 mL trypsin, 70 mL 10% DMEM, and fibrinogen stock solution to 37°C for 20 min.
3.4 Prepare the cell suspensions
3.4.1 Wash LX2 and HepG2 cells in T175 culture flasks twice with 10 mL PBS.
3.4.2 Add 6 mL trypsin to each flask and incubate for 4 min at 37 °C to allow cells to detach.
3.4.3 Once cells have detached add 6 mL 10% DMEM to each flask and inactivate trypsin.
3.4.4 Collect the cell suspension in 15ml tube and centrifuge for 3 min at 300 x *g*.
3.4.5 After centrifugation aspirate the supernatant and suspend each cell line in 5 mL 10% DMEM.
3.4.6 Cells are now counted using an automated cell counter, 10 µL of each cell suspension is added to the counting chamber slide and the slide is inserted into the cell counter. Cell count is displayed as cells/mL.
3.4.7 Using the cell counts from the cell counter the cells from each cell line can now be diluted to 1 × 10^6^ cells per mL into clearly marked 15 mL tubes.
3.4.8 Centrifuge the dilutions for 3 min at 300 x *g*
3.5 After centrifugation the supernatant is aspirated and 10% DMEM is added to each 15 mL tube according to the table below, values provided in the table is for 2 mL of each formulation.

3.6 Neutralize the amount of collagen with 10 µl/mL NaOH, and add the neutralized collagen to the cell suspension, the 10% DMEM present will turn yellow, once the suspension is mixed thoroughly by pipetting with a cut tip it will turn a bright pink.
3.7 The fibrinogen is now added to the collagen cell suspension according to the table, using a cut pipette tip, mix the suspension thoroughly.
3.8 Finally add thrombin to collagen-fibrinogen cell suspension, 0.1 KIU thrombin for each 10 mg of fibrinogen.
3.9 Remove the inverted inserts from the incubator and using a cut 200 µL pipette tip, pipette 200 µL of the prepared suspension onto the designated inserts.
3.10 Allow the gels to crosslink for 15 min within the laminar flow hood.
3.11 After 15 min, gently place the bottom section of the plate over the gels and move them to the incubator to allow the gels to crosslink at 37 °C for 45 min.
3.12 Once the gels have completed crosslinkage the inserts can be inverted again and 2 mL 10% DMEM can be added to each of the bottom wells of the plate.

| Formulation | Fibrinogen (mL) | Collagen (mL) | 10 % DMEM (mL) | Thrombin (µL) |
| --- | --- | --- | --- | --- |
| 1 | 1 | 0.8 | 0.2 | 4 |
| 2 | 0.75 | 0.8 | 0.45 | 3 |
| 3 | 0.25 | 0.8 | 0.95 | 1 |

### 4 Seeding endothelial cells

4.2 In a water bath, preheat 25 mL hanks balanced salt solution (HBSS), 10 mL trypsin, 10 mL trypsin inhibitor and 50 mL endothelial growth medium, to 37°C for 20 min.
4.3 Prepare the HUVEC cell suspension.
4.3.1 Wash HUVEC cells in T175 culture flasks twice with 10 mL HBSS.
4.3.2 Add 6 mL trypsin to the flask and incubate for 4 min at 37 °C to allow cells to detach.
4.3.3 Once cells have detached add 6 mL trypsin inhibitor to each flask to inactivate trypsin.
4.3.4 Collect the cell suspension in 15ml tube and centrifuge for 3 min at 200 x *g*.
4.3.5 After centrifugation aspirate the supernatant and suspend the cells in 5 mL endothelial growth medium.
4.3.6 Count the cells as previously described using an automated cell counter.
4.3.7 Using the cell count from the cell counter, prepare seeding density of 1.0 ×10^4^ cells/mL
4.4 Seed 500 µL per well HUVEC cells into the top part of the insert.

### 5 Maintenance

5.1 Medium is exchanged every second day, spent culture medium is aspirated from both the well and the insert and fresh culture medium is added. A volume of 2 mL is added to the well containing the gel and 0.5 mL to the insert. The model is maintained for 21 days prior to experimentation.

### 6 Rheology

6.1 Measure storage moduli of gel formulations to indicate stiffness values using a rheometer, by performing frequency sweeps from 0.1-20 Hz at 0.267% and 37°C, with a constant axial force of 0.1N using an 8 mm diameter parallel plate stainless steel geometry.

### 7 Viability and drug response

7.1 Drug response and viability was determined in 2D co-cultures and our 3D model.
7.1.1 Cells for the 2D co-culture were seeded onto black clear bottom 96-well plates at a seeding density of 5.0 × 10^4^ cells/mL. HepG2 and LX2 cells were seeded in a 1:1 ratio. Cells were allowed to attach overnight prior to doxorubicin treatment.
7.1.2 Our biomimetic model was setup and maintained for a period of 21-days.
7.2 Two hours prior to doxorubicin treatment, culture medium was aspirated from both the cells and the biomimetic model. Both models were washed twice with PBS. Starvation medium was added to the 2D co-culture (200 µL per well) and to the 3D model (2 mL to the wells containing the hydrogel and 500 µL to the insert).
7.3 Doxorubicin was administered to the both the 2D cell culture and the 3D constructs at dosages 0.5, 1 and 1.5 mM corresponding to the IC25, 50 and 75 values respectively as determined in 2D HepG2 cultures over 72h. Both models were treated with doxorubicin for 72h.
7.4 After 72h, the culture medium was aspirated from both the 2D cells and 3D constructs. Culture medium for the wells containing cells alone or imbedded into hydrogels were aspirated and the wells were wash three times with PBS.
7.5 AlamarBlue^®^ was prepared according to the manufacturer recommendations. The AlamarBlue^®^ reagent (150 µL per well for 2D cultures, 2 mL per well and 500 µL per insert for 3D culture) was added and the plate was incubated overnight at 37°C.
7.6 Following incubation 150 µL was transferred from each well of the 3D setup into a black clear bottom 96-well plate, the 2D co-culture was read within the plate they were seeded in.
7.7 Fluorescence was read using a micro plate reader with excitation wavelength at 485 and emission wavelength at 550 nm.
7.8 Results were calculate using the following formula:

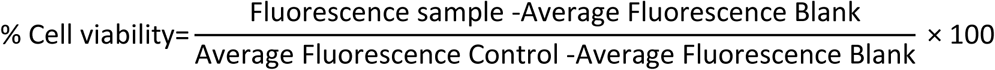

## REPRESENTATIVE RESULTS

### Protocol optimization

#### Concentration ranges and seeding volume

Protocol optimization occurred according to the schematic diagram presented in figure 2. Two physiologically relevant hydrogels, Collagen type I and Fibrinogen, was identified by means of a literature search. Starting with rat tail collagen type I, a range of concentrations (4, 3, 2 and 1 mg/mL) was seeded onto inverted inserts to determine their ability to successfully adhere to the insert once inverted again. All concentrations within this range were able to form gels, however collagen gels appeared flattened and had various air bubbles trapped within them as a result of handling and pipetting, see figures 3 and 4. To determine the optimal seeding volume in an attempt to improve the quality of the collagen gel, a range of seeding volumes (100, 150 and 200 µL), was seeded onto inverted inserts, see figure 4. The seeding volume had no influence on the appearance of the gel or the presence of bubbles within the gel. Accordingly it was decided that 200 µL was the optimal seeding volume producing the fullest gels. Fibrinogen from bovine was also evaluated for its ability to produce a gel that could adhere to an insert for a prolonged time period. A concentration range (70, 50, 40, 30, 20, 10, 5, 1 mg/mL) was prepared and seeded at a volume of 200 µL onto the lid of a 12-well plate, see figure 5. Within 20 min after seeding all the concentrations were able to successfully form a gel. However, the concentrations 5 and 1 mg/mL was excluded as the gels formed had a fluid like consistency and had started to detach from the lid after being kept overnight at 37 °C.

**Figure 1:**
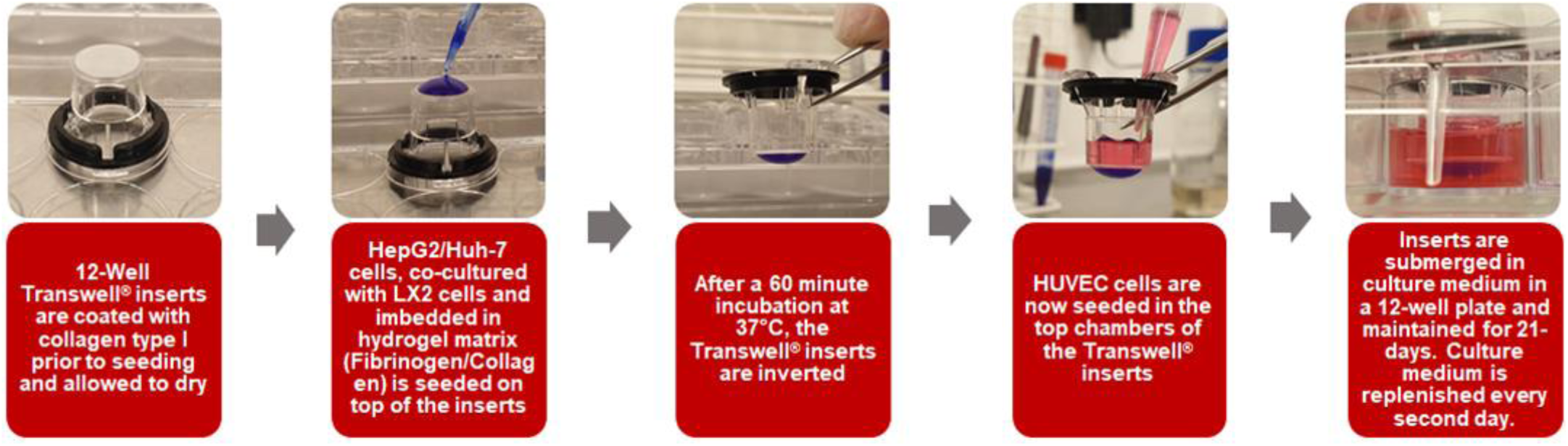
Graphical depictions of the creation of the 3D HCC Transwell™ model.

**Figure 2:**
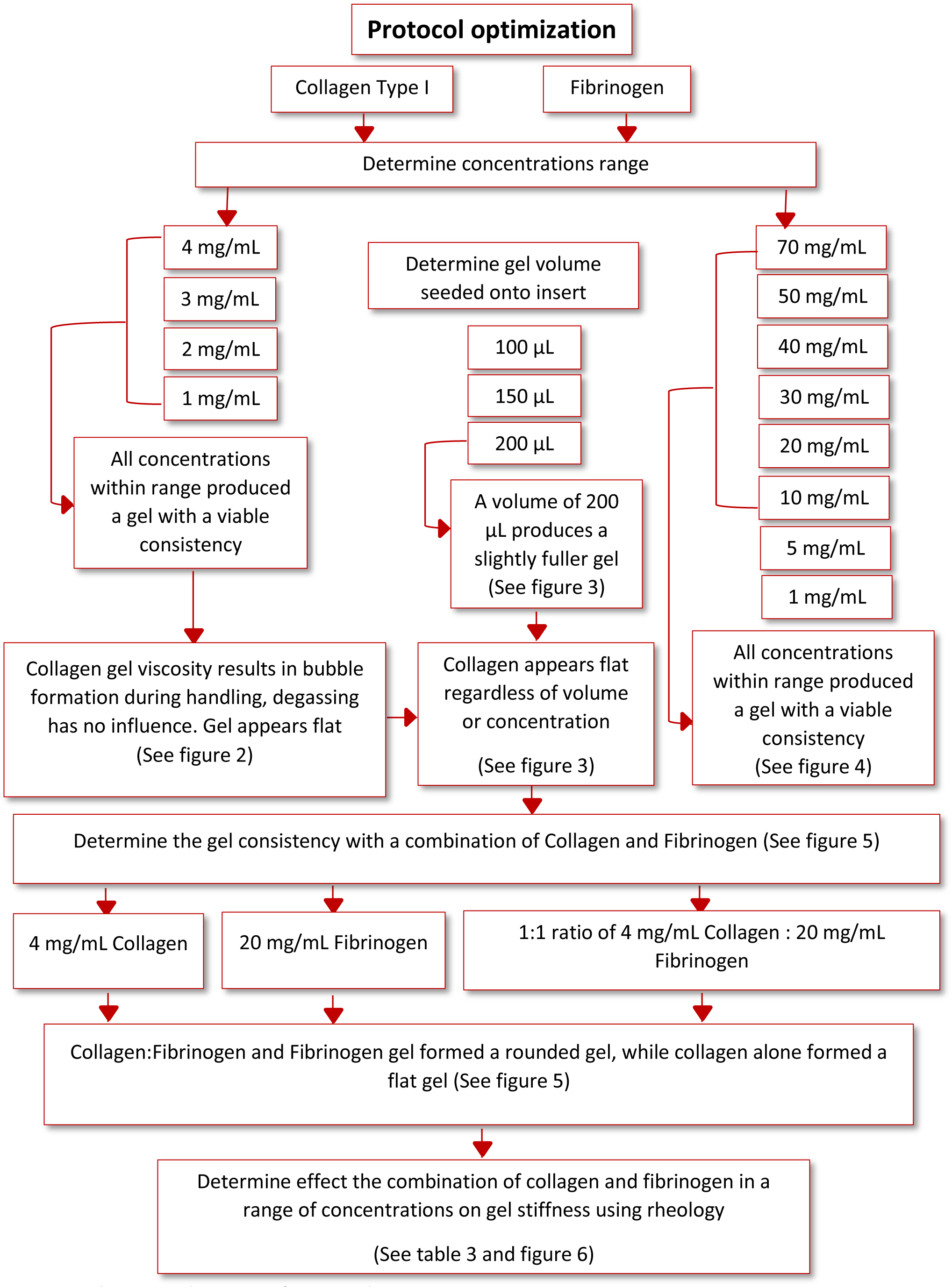
Schematic diagram of protocol optimization.

**Figure 3:**
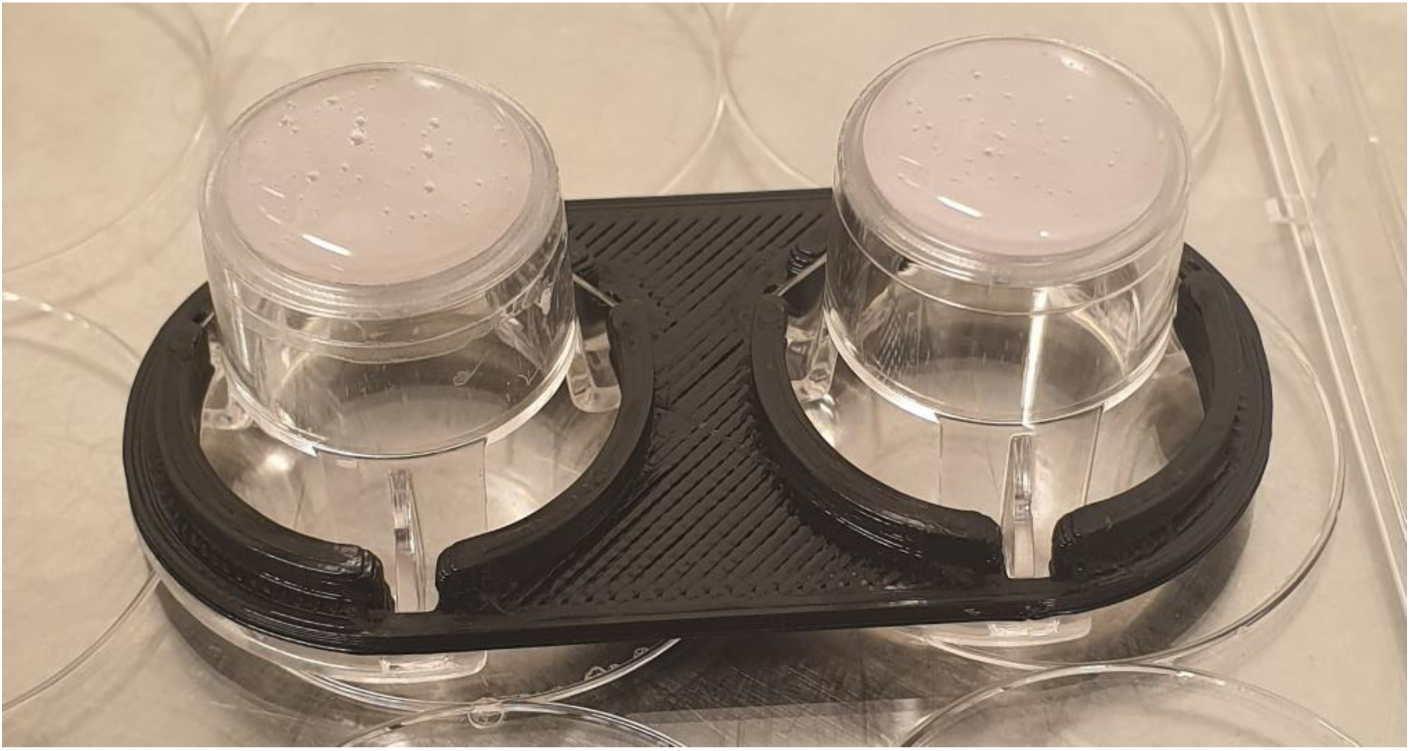
Collagen gels 4 mg/mL containing 1.0 × 10^5^ cells/mL seeded onto inserts showing bubbles present in the gel. Insert on the left been degassed on ice for 15 min, while insert on the right has not been degassed. Degassing seems to have no effect on the bubbles present in the gels

**Figure 4:**
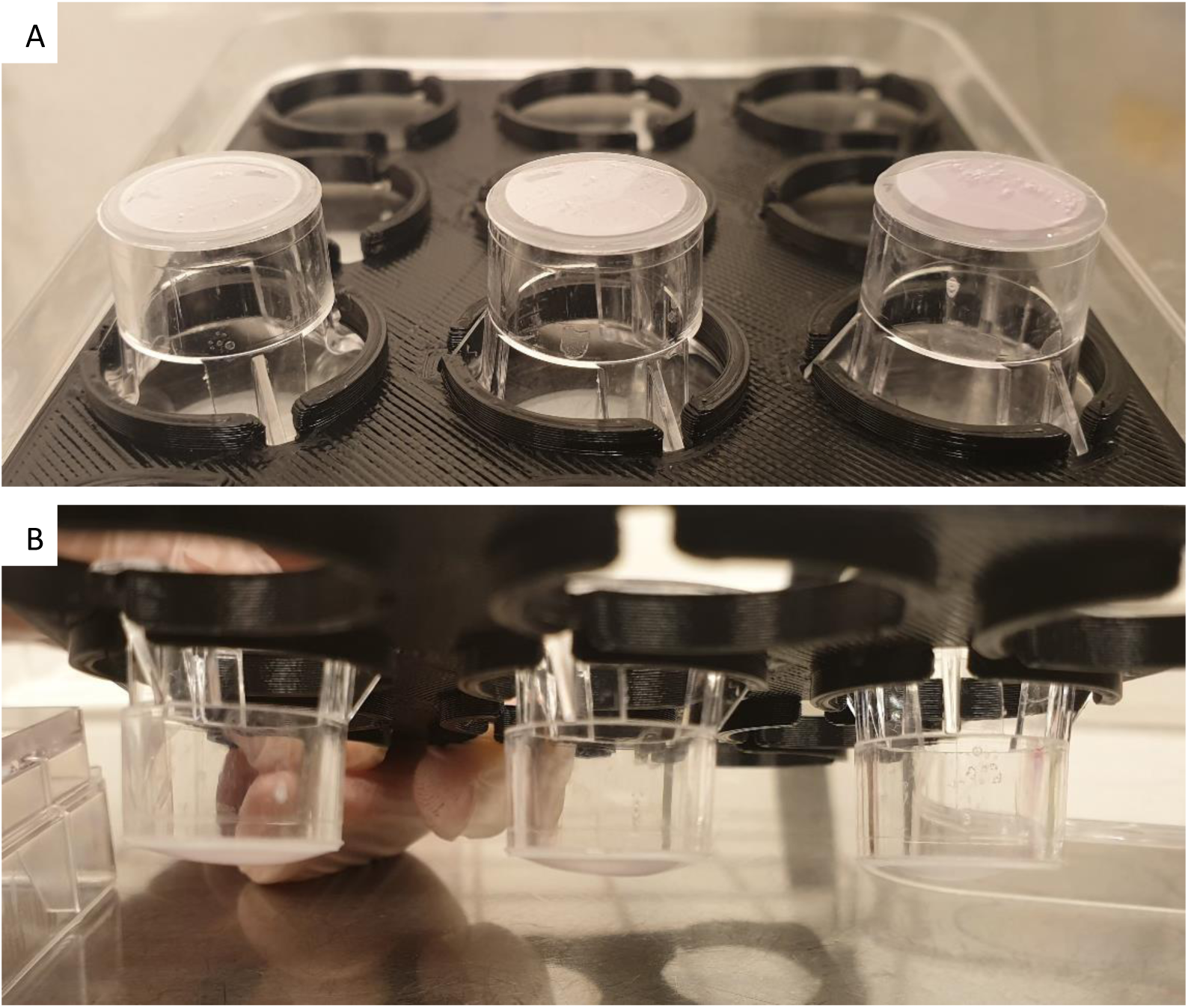
Collagen gels 4 mg/mL containing 1.0 × 10^5^ cells/mL seeded onto inserts in varying volumes. **(A)** Insert on the left 100 µL, middle 150 µL and right 200 µL, directly after seeding. Bubbles in the gels are still present. **(B)** Insert on the left 200 µL, middle 150 µL and right 100 µL, after 60 min crosslinking. All gels regardless of volume still appear flat.

**Figure 5:**
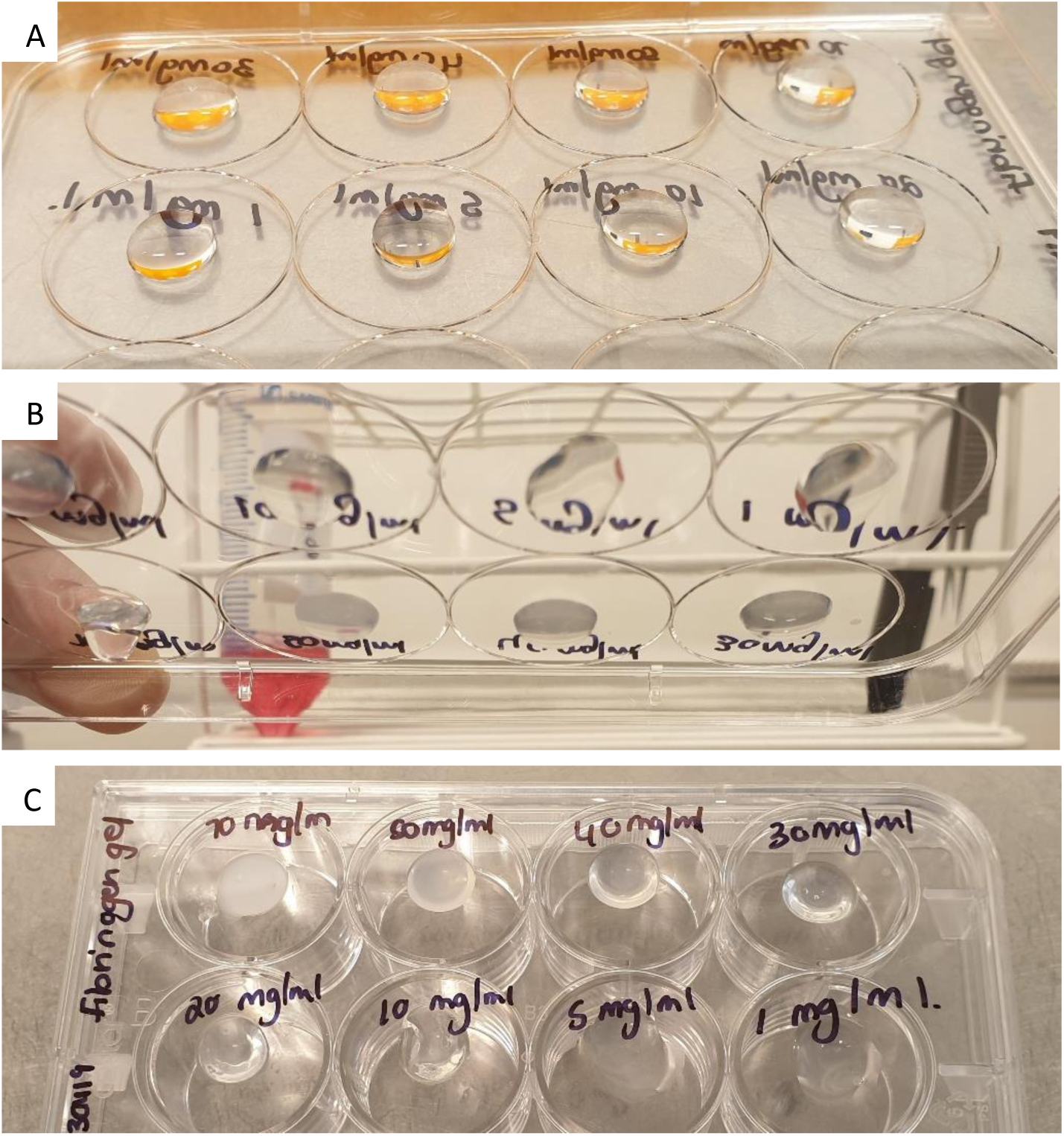
Fibrinogen (200 µL) in a concentration range from 70 to 1 mg/mL seeded onto the lid of a 12-well plate. **(A)** Gels directly after seeding. **(B)** Gels 20 min after seeding. **(C)** Gels kept overnight. All gels appear well rounded, the 5 mg/mL and 1 mg/mL gels were excluded as they appeared to have a more fluid consistency 20 min after seeding and had started to detach from the lid after being kept overnight.

#### Combination collagen and Fibrinogen

Based on the results from the collagen and fibrinogen concentration ranges the effect of combining the collagen and fibrinogen was evaluated. Three inserts were setup with the following, 4 mg/mL collagen, 20 mg/mL fibrinogen and a 1:1 ration of collagen (4 mg/mL) and fibrinogen (20 mg/mL), see figure 6. Directly after seeding the hydrogels, we observed that the insert with collagen alone still had a flat appearance combined with the occurrence of bubbles within the gel. The inserts with fibrinogen produced a full and rounded gel, so did the insert with the combination of collagen and fibrinogen. After cross linking at 37 °C for 60 min, all gels had attached to the insert and remained attached following overnight incubation at 37 °C.

**Figure 6:**
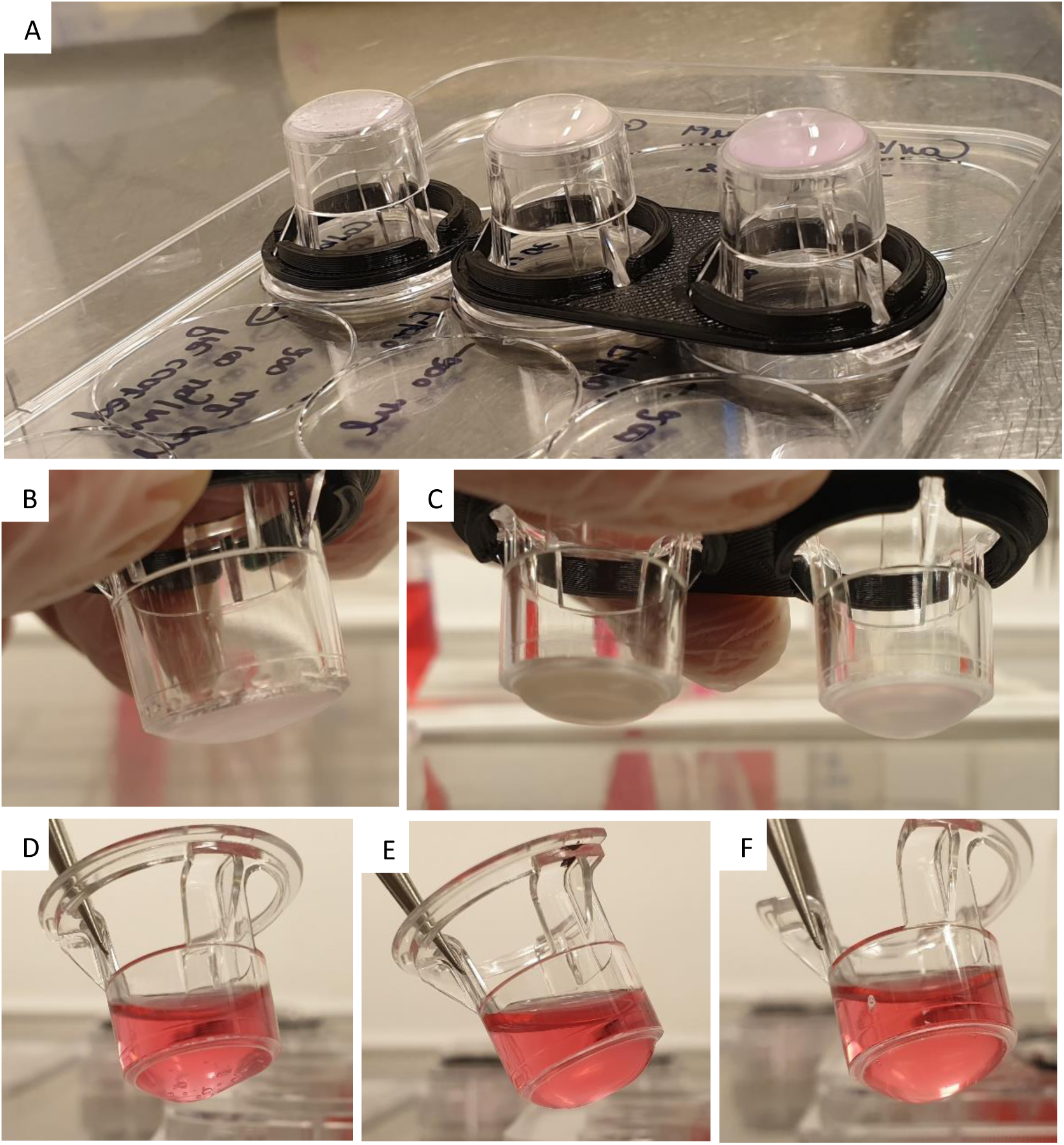
**(A)** Collagen 4 mg/mL seeded onto insert on the left, fibrinogen 20 mg/mL seeded onto insert in the middle and collagen 4 mg/mL, fibrinogen 20 mg/mL (1:1 ration) seeded onto insert on the right. All gels seeded at a volume of 200 µL containing 1.0 × 10^5^ cells/mL, all gels were crosslinked for 60 min at 37 °C. **(B)** Collagen 4 mg/mL 60 min after crosslinking, gel appears flat and contains bubbles. **(C)** Insert on the left fibrinogen 20 mg/mL, gel appear rounded, no bubbles present, insert on the right collagen 4 mg/mL, fibrinogen 20 mg/mL gel, gel is well rounded with no bubbles. **(D)** Collagen 4 mg/mL gel after being kept overnight at 37 °C, gel still contains a large amount of bubbles, some swelling of the gel has occurred. **(E)** Fibrinogen 20 mg/mL kept overnight at 37 °C. **(F)** Collagen 4 mg/mL, fibrinogen 20 mg/mL gel kept overnight at 37 °C.

#### Determining Cell seeding density

Based on previous experience working with hydrogels an experiment was setup to determine the optimal cell seeding density (data not shown). Cells were embedded into a combination of collagen and fibrinogen in the following concentration range (7.5 × 10^5^, 8.5 × 10^5^, 9.5 × 10^5^, 1.0 × 10^6^, 1.5 × 10^6^ and 2.0 × 10^6^ cells/mL), it was found that 2.0 × 10^6^ cells/mL as in Smit et al., 2020, was the optimal seeding density.

#### Rheology

Ten formulations of the fibrinogen and collagen hydrogel combinations were evaluated by means of rheology, see table 3. The aim was to determine which of these formulations could mimic liver stiffness seen during the development of HCC. Literature provided known liver stiffness values for rats, mice and humans during fibrosis, cirrhosis and HCC and the aim was to get as close as possible to these values. The ten formulations as set out in table 3 was prepared in triplicate and the storage modulus of each were determined using a rheometer, results not shown.

**Table 3:** Collagen and fibrinogen combinations in various concentrations evaluated with Rheology to determine stiffness values

| Formulation | Fibrinogen (mg/ml) | Collagen (mg/ml) |
| --- | --- | --- |
| 1 | 60 | 2 |
| 2 | 50 | 2 |
| 3 | 40 | 2 |
| 4 | 30 | 2 |
| 5 | 20 | 2 |
| 6 | 10 | 2 |
| 7 | 20 | 5 |
| 8 | 20 | 4 |
| 9 | 20 | 3 |
| 10 | 20 | 1 |

From these ten formulas three was chosen to proceed with. These included 2 mg/mL collagen type I and 10 mg/mL fibrinogen corresponding to liver stiffness values at the onset of fibrosis, 2 mg/mL collagen type I and 30 mg/mL fibrinogen corresponding to cirrhosis and 2 mg/mL collagen type I and 40 mg/mL fibrinogen corresponding to HCC, see figure 7. The data also indicated that varying collagen values do not result in notable changes of stiffness values when compared to varying values of the fibrinogen.

**Figure 7:**
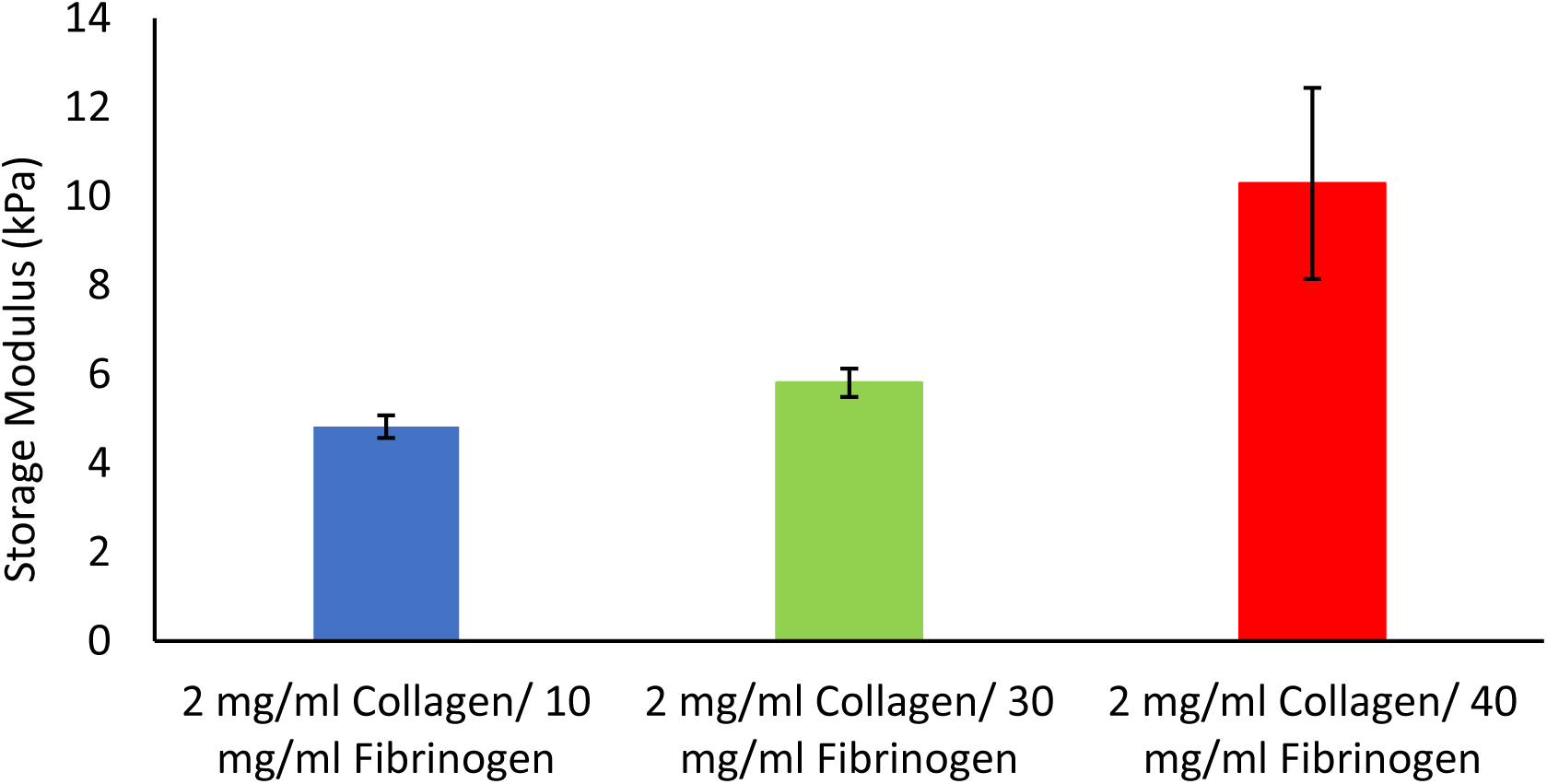
Hydrogel stiffness values for various Fibrinogen/Collagen hydrogel formulations measured. Storage modulus and loss modulus were determined at 37°C and 1Hz by means of a Discovery HR-2 Hybrid Rheometer (n = 3, Error bars = SD)

#### Viability, drug response and metastatic potential

The results from the AlamarBlue^®^ assay showed an overall reduced cell viability within the 2D co-culture, lower than expected based on the known reported IC 25, 50 and 75 values, when compared to the untreated control, see figure 8. This may be attributed to the LX2 cells in our co-culture that are more sensitive to Doxorubicin treatment. However, in our 3D model we noticed and increase in doxorubicin resistance, confirming the decrease in chemotherapeutic potential often seen in 3D model systems. Statistical significance compared to controls was assessed using the Student T test (two-tailed), with P<0.05 considered significant.

**Figure 8:**
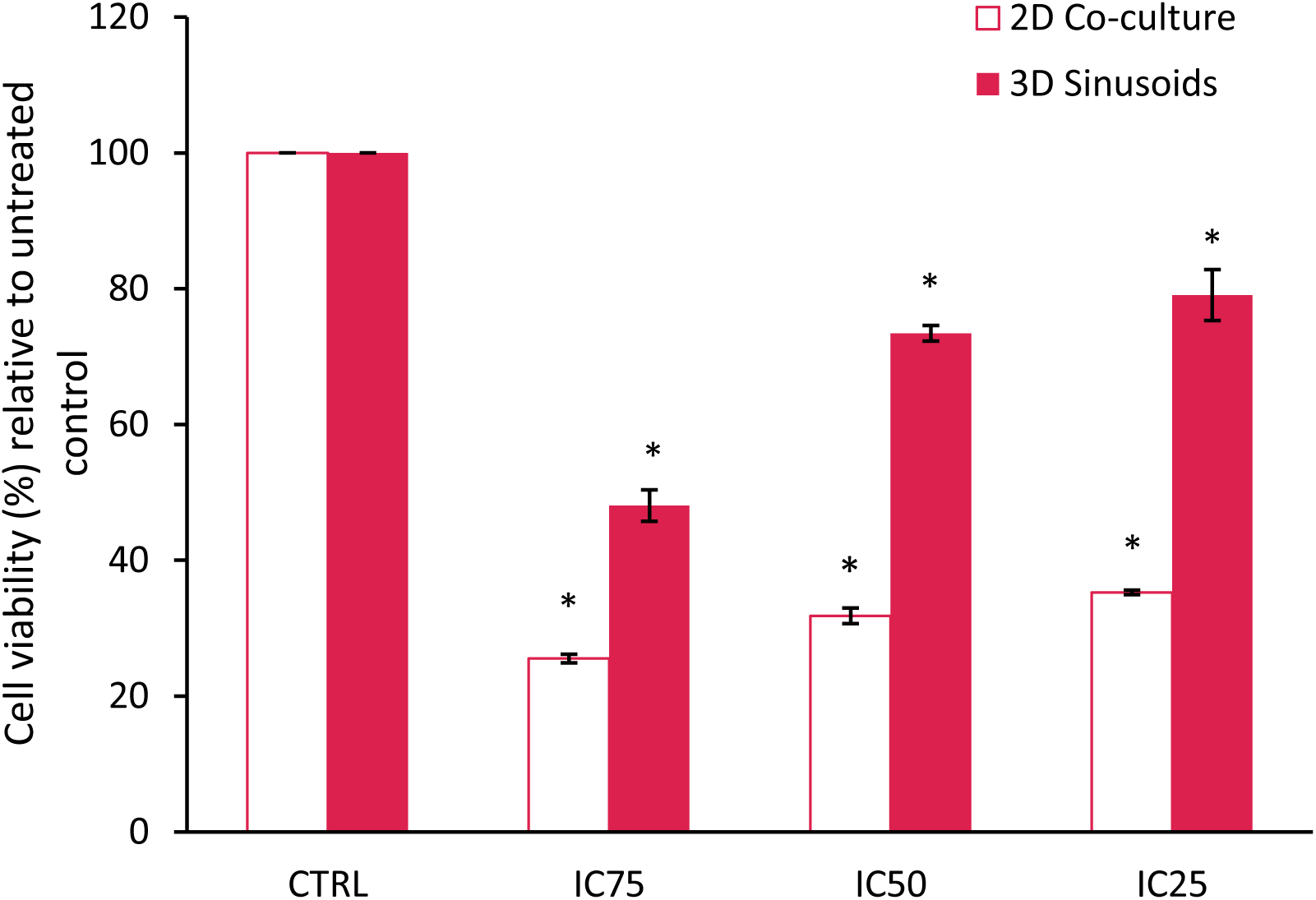
Percentage cell viability of a 2D co-culture model compared to a 3D sinusoid co-culture model after treatment with Doxorubicin at various concentrations for a period of 72h. Results normalized relative to the untreated control (n=3, error bars = SD) (* = p<0.0001).

The gel constructs were visually inspected daily using a light microscope to follow cell growth within different concentrations of the hydrogels. Cells filled up the hydrogels in a homogeneous and compact way, from day 7 spheroids started to assemble within the matrix. From day 17 onwards the model started to became metastatic, see figure 9. Samples displayed clusters of spheroids and cells dissociating from the gel, floating in the medium. This unexpected finding lead us to believe that our model may also serve as a future metastatic model.

**Figure 9:**
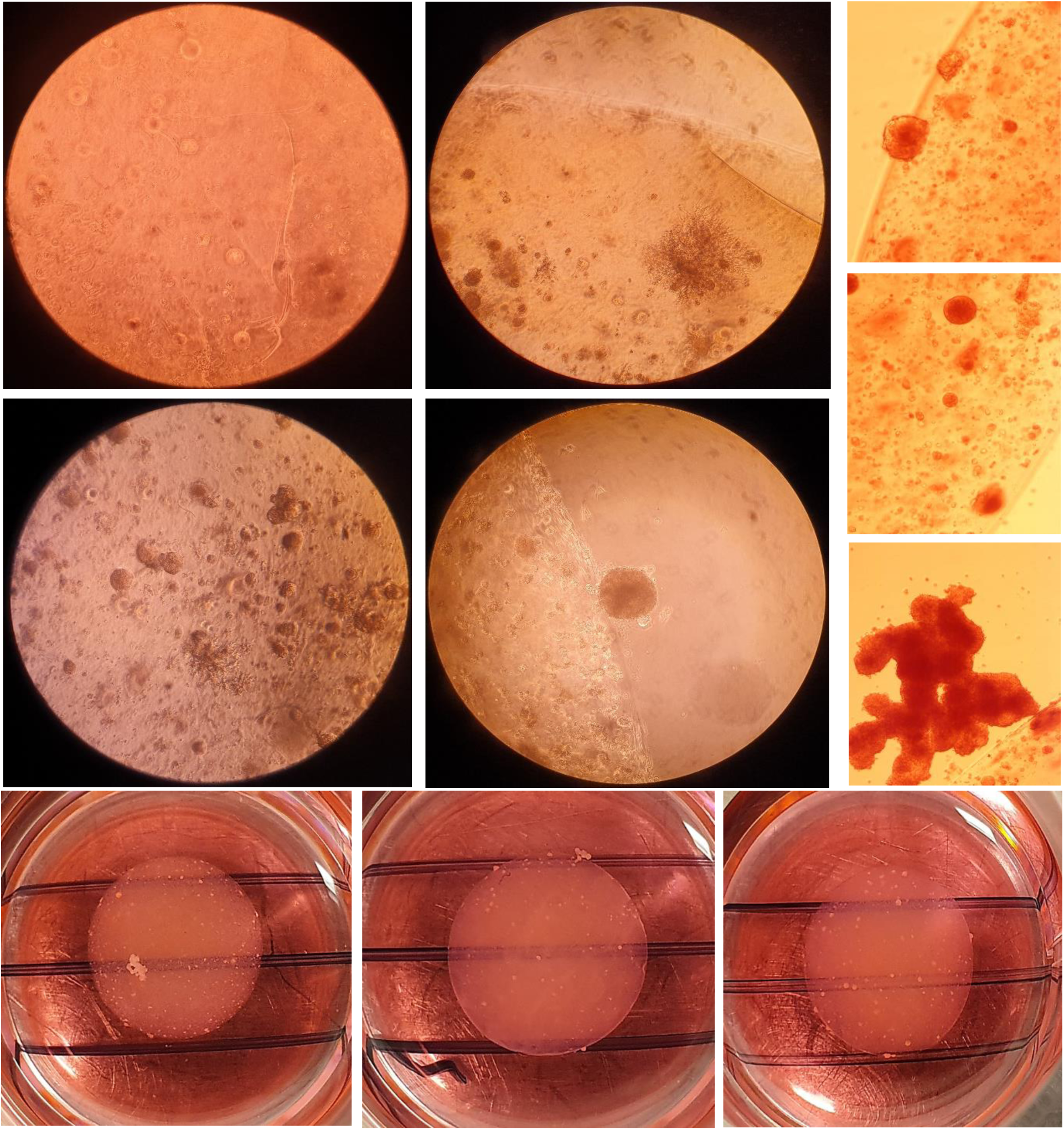
Metastatic tumor nodules formation on the edges and within of hydrogel constructs

## DISCUSSION

This protocol describes the development of a method to create a biomimetic HCC model. A clear workflow has been established and the critical steps involved identified. These include, preparation of the fibrinogen stock solution, coating the inserts with collagen and seeding the cells imbedded into the hydrogel. During the preparation of the fibrinogen stock solution it is important to add the fibrinogen in smaller increments at higher concentrations. This will not only reduce the time it takes for the fibrinogen to dissolve, but will also prevent the fibrinogen from gelling inconsistently and prematurely as seen in figure 10. The preparation of the fibrinogen gel takes a considerable amount of time and this may influence the overall experimental success. Results indicates that once the fibrinogen gel starts to gel inconsistently it is best to discard it. The inserts should be coated with collagen and washed with PBS and dried within the laminar flow hood prior to seeding the cells embedded in hydrogels. Failure to ensure that the inserts are dry will result in the hydrogels spilling over the edges of the insert that will result in an uneven gel. Unevenness of the gel will ultimately influence results where diffusion is a factor.

**Figure 10:**
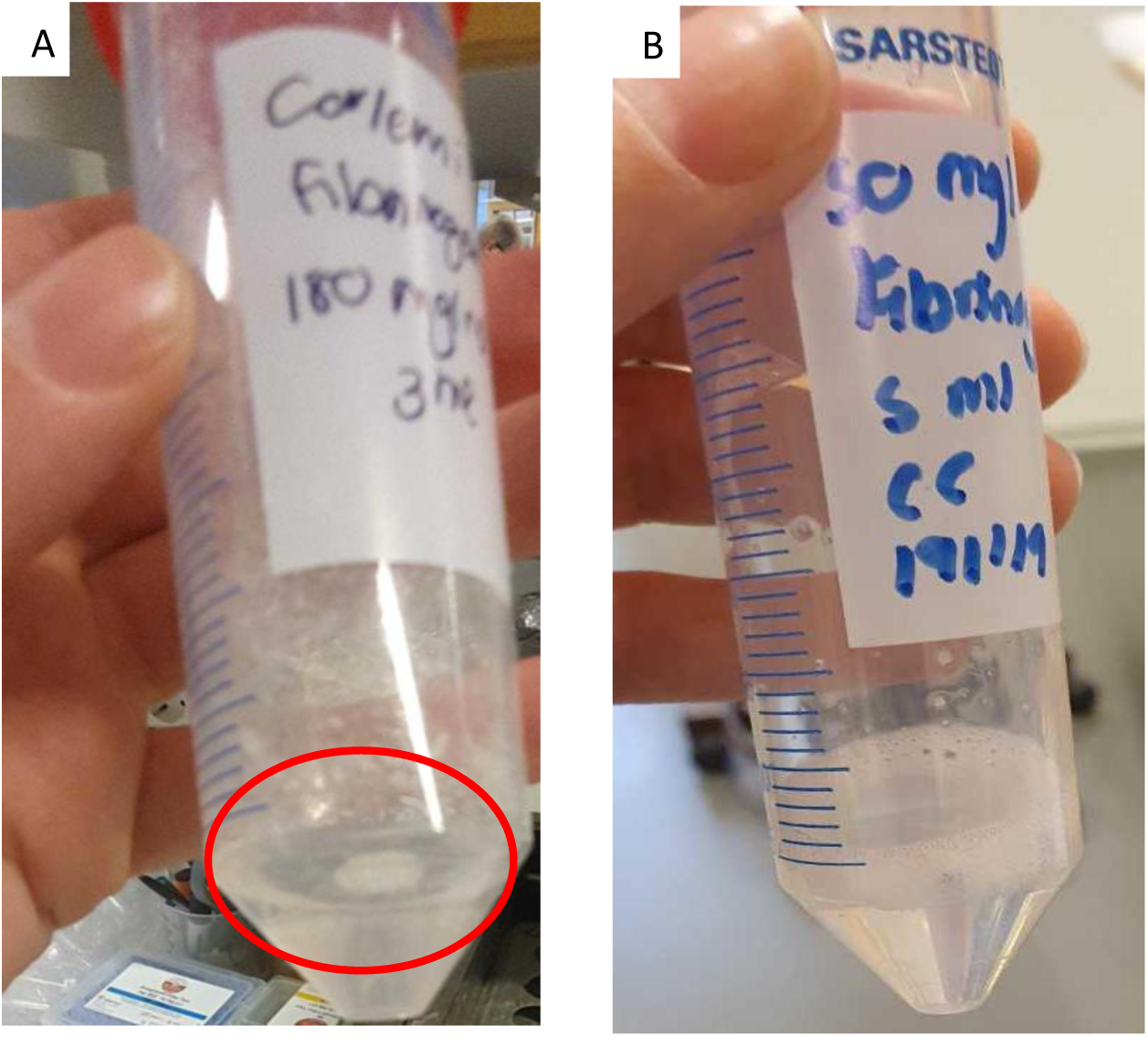
Preparation of fibrinogen gel for Fibrinogen/Collagen hydrogel formulations. **(A)** Fibrinogen gel solution that has formed clumps and started to gel prematurely with undissolved fibrinogen adhering to the tube. **(B)** Fibrinogen gel solution that has dissolved completely, solution is clear and slightly more viscous.

It is recommended to work as fast as possible while seeding the hydrogel cell suspension onto the inserts, as the fibrinogen component will start to crosslink with the addition of thrombin. Prepare smaller working volumes at a time when working with gel suspensions at higher concentrations to prevent the gel from crosslinking while seeding as this will have an effect on the distribution and amount of gel seeded onto each well. The order of adding the components is critical, in this protocol we provided a streamlined workflow to prevent gels from crosslinking prematurely. Due to the viscosity of the hydrogel gel suspension working with a cut pipette tip is advised during mixing and measuring. When mixing the suspension ensure that this is done quickly and evenly to create a homogenous suspension. Uneven mixing will result in a heterogeneous gel which will negatively affect results, see figure 11.

**Figure 11:**
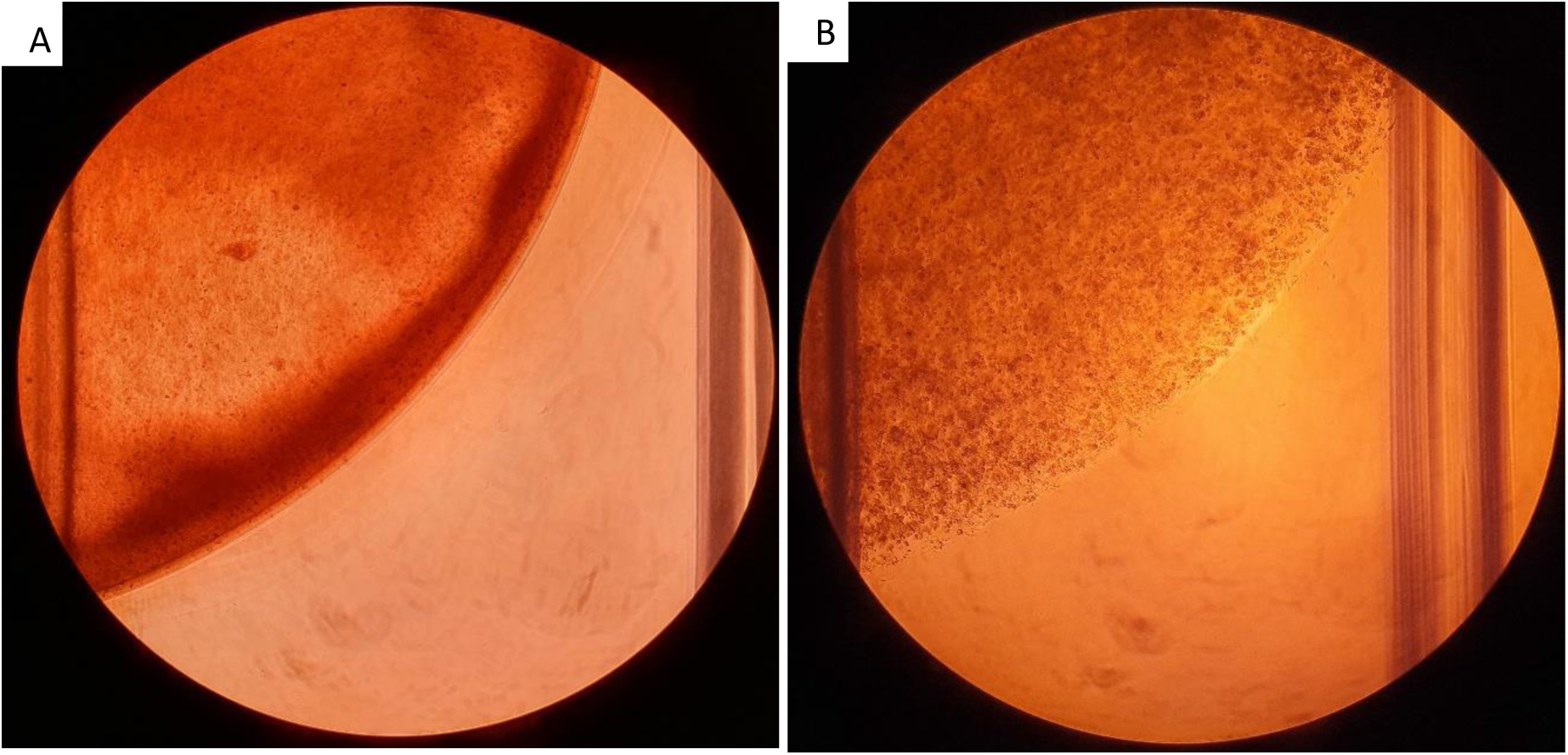
Collagen/Fibrinogen gels seeded onto 12-well plates, all gels seeded at a volume of 200 µL containing 2mg/mL Collagen and 20 mg/mL Fibrinogen with 2.0 × 10^6^ cells/mL. **(A)** Hydrogel gel with heterogeneous consistency, visible uneven distribution of the hydrogels. **(B)** Hydrogels homogenously mixed.

Following the protocol optimization the model was evaluated to determine the models bio-physical properties. Rheology data showed that our model, composed of physiologically relevant extracellular matrix components namely collagen type I and fibrinogen, was able to mimicking the bio-physical properties of a fibrotic, cirrhotic and HCC liver. Recapitulating liver stiffness in 3D models for HCC is of considerable importance and is often overlooked during model development. Increased liver stiffness is related to chemotherapeutic resistance, proliferation, migration and dormancy within in HCC (Schrader *et al*., 2011). While the activation of hepatic stellate cells in HCC is associated with increased extracellular matrix rigidity, with several signaling pathways associated with these hepatic stellate cells showing mechanosensitivity (Lachowski *et al*., 2019).

The inclusion of stroma associated cells in the development of 3D models for HCC has become increasingly relevant. Studies show that multicellular spheroids composed of hepatic stellate and HCC cells resulted in increased chemotherapeutic resistance and invasive migration, while mimicking HCC tumor appearance *in vivo*, when compared to a PXT mice model and human HCC tissue samples (Khawar *et al*., 2018). A similar study by Jung *et al*., 2017 found multicellular spheroid consisting of Huh-7 and HUVEC cells promoted vascularization and aggressiveness. These spheroids showed viability at significantly higher concentrations of doxorubicin and sorafinib when compared to Huh-7 monoculture spheroids. During the evaluation of our models viability and response to doxorubicin, we found that our model, with varying stiffness values and the inclusion of stroma associated cells, showed a similar decreased in response to chemotherapeutics when compared to a 2D co-culture model. Thus effectively mimicking drug resistance typically seen in patients and other 3D HCC models. In addition, our model also gave rise to metastatic tumor nodules, providing an interesting new platform to study multifocal HCC or to identify mechanisms that contribute to early stages of metastasis.

As this is a modular system the model can be fortified by the addition of other extracellular matrix components namely, laminin and hyaluronic acid. Alternatively the current hydrogels used within this model can be replace by synthetic hydrogels such as sodium alginate or chitosan. Further modifications to the current model is the substitution of the cell lines with primary cell cultures to produce an even more physiologically relevant model.

We have thus successfully developed a 3D model with tunable bio-physical properties for studying tumor-stroma interactions in HCC. We have found our model to be more representative of the *in vivo* situation when compared to traditional 2D cultures. However, there is still much to be done, we hope to extensively characterize this model and explore the model as a possible metastatic platform to answer more complex and pressing questions that remain in the study HCC.

## ACKNOWLEDGMENTS

This research was funded through grants obtained from the Swedish Cancer Foundation (Cancerfonden, CAN2017/518), the Swedish society for medical research (SSMF, S17-0092), the O.E. och Edla Johanssons foundation and the Olga Jönssons foundation. These funding sources were not involved in the study design; collection, analysis and interpretation of data; writing of the report; and in the decision to submit the article for publication. 3D printing of custom designed spacers used in this protocol was performed at U-PRINT: Uppsala University’s 3D-printing facility at the Disciplinary Domain of Medicine and Pharmacy. We would like to thank Paul O’Callaghan for his valuable input on our project.

## REFERENCES

Galle, P.R., Forner, A., Llovet, J.M., Mazzaferro, V., Piscaglia, F., Raoul, J-L., Schrmacher, P. and Vilgrain, V. 2018. EASL Clinical practice guidelines: Management of hepatocellular carcinoma. Journal of Hepatology., 69:182–236.

Marquardt, J.U., Andersen, J.B. and Thorgeirsson, S.S. 2015. Functional and genetic deconstruction of the cellular origin in liver cancer. Nature Reviews Cancer., 15:653–667.

Balogh, J., Ill, D.V., Asham, E.h., Burroughs, S.G., Boktour, M., Saharia, A., Li, X., Ghobrial, R.M. and Monour, H.P. 2016. Hepatocellular Carcinoma: A review. Journal of Hepatocellular Carcinoma., 3:41–53.

Perumpail, R.B., Womg, R.J., Ahmed, A. and Harrison, S.A. 2015. Hepatocellular carcinoma in the setting of non-cirrhotic non-alcoholic fatty liver disease and the metabolic syndrome: US experience. Dig Dis Sci., 60:3142–3148.

Baglieri, J., Brenner, D.A. and Kisseleva, T. 2019. The role of fibrosis and liver associated fibroblasts in the pathogenesis of hepatocellular carcinoma. International Journal of Molecular Sciences., 20:1723 doi: 10.3390/ijms20071723

Arriazu, E., De Galarreta, M.R., Cubero, F.J., Varela-Rey, M., De Obanos, M.P.P., Leung, T.M., Lopategi, A., Benedicto, A., Abraham-Enachescu, I. and Nieto, N. 2014. Extracellular matrix and liver disease. Antioxidants & Redox Signaling., 21(7): 1078–1097. doi: 10.1089/ars.2013.5697

Malarkey, D.E., Johnson, K., Ryan, L., Boorman, G. and Maronpot, R.R. 2005. New insight into functional aspects of liver morphology. Toxicologic PathologyI., 33:27–34. doi: 10.1080/01926230590881826

Moreira, R.K. 2007. Hepatic stellate cells and liver fibrosis. Arch Pathol Lab Med., 131: 1728–1734.

Hernandez-Gea, V., Toffanin, S., Friedman, S.L and Llovet, J.M. 2013. Role of the microenvironment in the pathogenesis and treatment of hepatocellular carcinoma. Gastroenteroloy 144:512–527.

Amicone, L. and Marchetti, A. 2018. Microenvironment and tumor cells: two targets for new molecular therapies of hepatocellular carcinoma. Translational Gastroenterology and Hepatology., 3:24. http://dx.doi.org/10.21037/tgh.2018.04.05

Rawal, P., Siddiqui, H., Hassan, M., Choudhary, M.C., Tripathi, D.M., Nian, V., Trehanpati, N. and Kaur, S. 2019. Endothelial cell-derived TGF-B promotes epithelial-mesenchymal transition via CD133 in Hbx-Infected Hepatoma cells. Frontiers in Oncology., 9(308):1–9 doi.org/10.3389/fonc.2019.00308

Yoo, J.E., Kim, Y-J., Rhee, H., Kim, H., Ahn, E.Y., Choi, J.S., Roncalli, M. and Park, Y.N. 2017. Progressive enrichment of stemness features and tumour stromal alterations in multistep hepatocarcinogenesis. PlosONE., doi: 10.1371/journal.pone.0170465.g001.

Landry, B/D., Leete, T., Richards, R., Cruz-Gordilla, P., Schwartz, H.R., Honeywell, M.E., Ren, G., Schwartz, A.D., Peyton, S.R and Lee, M.J. 2018. Tumor-stroma interactions differentially alter drug sensitivity based on the origin of stromal cells. Molecular systems biology., 14:e8332 doi: 10.15252/msb.20188322.

Le, BD, Kang, D, Yun, S, Jeong, YH, Kwak, J-Y, Yoon, S, and Jin, S. 2018. Three-dimensional hepatocellular carcinoma/fibroblast model on a nanofibrous membrane mimics tumor cell phenotypic changes and anticancer drug resistance. Nanomaterials., 8(64): 1–11.

Lv, D., Hu, Z., Lu, L., Lu, H. and Xu, X. 2017. Three-dimensional cell culture: A powerful tool in tumor research and drug discovery (Review). Oncology Letters., 14: 6999–7010.

Hoarau-Véchot J, Rafii A, Touboul, C and Pasquier, J. 2018. Halfway between 2D and animal models: Are 3D cultures the ideal tool to study cancer-microenvironment interactions? International Journal of Molecular Sciences, 19(181), 1–24.

Khawar, I.A., Park, J.K., Jung, E.S., Lee, M.A., Chang, S., Kuh, H.-J., 2018. Three Dimensional Mixed-Cell Spheroids Mimic Stroma-Mediated Chemoresistance and Invasive Migration in hepatocellular carcinoma. Neoplasia 20, 800–812. https://doi.org/10.1016/j.neo.2018.05.008

Elliott, N.T. and Yuan, F. 2010. A review of three-dimensional *in vitro* tissue models for drug discovery and transport studies. Journal of Pharmaceutical Sciences., 100(1):59–74. DOI 10.1002/jps

Nath, S. and Devi, G. R. 2016. Three-dimensional culture systems in cancer research: focus on tumor spheroid model. Pharmacol Ther. 163:94–108. doi: 10.1016/j.pharmthera.2016.03.013.

Zanoni, M., Piccinini, F., Arienti, C., Zamagni, A., Santi, S., Polico, R., Bevilacqua, A. and Tesei, A. 2016. 3D tumor spheroid models for *in vitro* therapeutic screening: a systematic approach to enhance the biological relevance of data obtained. Scientific Reports., 6:19103 DOI: 10.1038/srep19103

Bell, C.C., Hendriks, D.F.G., Moro, S.M.L., Ellis, E., Walsh, J., Renblom, A., Puigvert, L.F., Dankers, A.C., Jacons, F., Snoeys, J., Sison-Young, R.L., Jenkins, R.E., Nordling, A., Mkrtchian, S., Park, B.K., Kitteringham, N.R., Goldrimg, C.E.P., Lauschke, V.M and Ingelman-Sundberg, M. 2016. Characterization of primary human hepatocyte spheroids as a model system for drug-induced liver injury, liver function and disease. Scientific Reports., 6:25187 DOI: 10.1038/srep25187

Jung, H-R., Kang, H.M., Ryu, J-W., Kim, D-S., Noh, K.H., Kim, E-S., Lee, H-J., Chung, K-S., Cho, H-S., Kim, N-S., Im, D-S., Lim, J.H. and Jung, C-R. 2017. Cell spheroids with enhanced aggressiveness to mimic human liver cancer *in vitro* and *in vivo*. Scientific Reports., 7:10499 DOI:10.1038/s41598-017-10828-7

Eilenberger, C., Rothbauer, M., Ehmoser, E-K., Ertl, P. and Küpcü, S. 2019. Effect of spheroidal age on sorafenib diffusivity and toxicity in a 3D hepg2 spheroid model. Scientific Reports., 9:4863 https://doi.org/10.1038/s41598-019-41273-3

K. Wrzesinski and S.J. Fey. 2013. After trypsinisation, 3D spheroids of C3A hepatocytes need 18 days to re-establish similar levels of key physiological functions to those seen in the liver. Toxicology Research; 2(2) 123–135. DOI: 10.1039/C2TX20060K

K. Wrzesinski, C.M. Magnone, L. Visby Hansen, M. Ehrhorn Kruse, T. Begauer, M. Bobadilla, M. Gubler, J. Mizrahi, C. Møller Andreasen, K. Zhang, K. Eyed Joensen, S.M. Andersen and S.J. Fey. 2013. Human liver spheroids exhibit stable physiological functionality for at least 24 days after recovering from trypsinisation. Toxicology Research; 2(3) 163–172 DOI: 10.1039/C3TX20086H

Mazza, G., Rombouts, K., Hall, A.R., Urbani, L., Luong, T.V., Al-Akkad, W., Longato, L., Brown, D., Maghsoudlou, P., Dhillon, A.P., Fuller, B., Davidson, B., Moore, K., Dhar, D., De Coppi, P., Malago, M. and Pinzani, M. 2015. Decellularized human liver as a natural 3D-scaffold of liver bioengineering and transplantation. Scientific Reports., 5:13079 DOI: 10.1038/srep13079

Ma, X., Yu, C., Wang, P., Xu, W., wan, X., Lai, C.S.E., Liu, J., Koroleva-Maharjh, A and Chen, S. 2018. Rapid 3D bioprinting of decellularized extracellular matrix with regionally varied mechanical properties and biomimetic microarchitecture.

Smit, T., Calitz, C., Willers, C., Svitina, H., Hamman, J., Fey, S.J., Gouws, C and Wrzesinski, K. 2020. Characterization of an alginated encapsulated LS180 spheroid model for anti-colorectal cancer compound screening. ACS Medicinal Chemistry Letters https://doi.org/10.1021/acsmedchemlett.0c00076

Schrader, J., Gordon-Walker, T.T., Aucott, R.L., Van Deemter, M., Quaas, A., Walsh, S., Benten, D., Forbes, S.J., Wells, R.G and Iredale, J.P. 2011. Matrix stiffness modulates proliferation, chemotherapeutic response, and dormancy in hepatocellular carcinoma cells. Hepatology 53(4) 1192–1205 doi: epdf/10.1002/hep.24108

Lachowski, D., Cortes, E., Rice, A., Pinato, D., Rombouts, K & Hernandez, A. 2018. Matrix stiffness modulates the activity of MMP-9 and TIMP-1 in hepatic stellate cells to perpetuate fibrosis. Scientific Reports., 9:7299 https://doi.org/10.1038/s41598-019-43759-6

